# The role of XPB/Ssl2 dsDNA translocation processivity in transcription-start-site scanning

**DOI:** 10.1101/2020.11.09.375329

**Authors:** Eric J. Tomko, Olivia Luyties, Jenna K. Rimel, Chi-Lin Tsai, Jill O. Fuss, James Fishburn, Steven Hahn, Susan E. Tsutakawa, Dylan J. Taatjes, Eric A. Galburt

## Abstract

The general transcription factor TFIIH contains three ATP-dependent catalytic activities. TFIIH functions in nucleotide excision repair primarily as a DNA helicase and in Pol II transcription initiation as a dsDNA translocase and protein kinase. During initiation, the XPB/Ssl2 subunit of TFIIH couples ATP hydrolysis to dsDNA translocation facilitating promoter opening and the kinase module phosphorylates the C-terminal domain to facilitate the transition to elongation. These functions are conserved between metazoans and yeast; however, yeast TFIIH also drives transcription start-site scanning in which Pol II scans downstream DNA to locate productive start-sites. The ten-subunit holo-TFIIH from *S. cerevisiae* has a processive dsDNA translocase activity required for scanning and a structural role in scanning has been ascribed to the three-subunit TFIIH kinase module. Here, we assess the dsDNA translocase activity of ten-subunit holo- and core-TFIIH complexes (i.e. seven subunits, lacking the kinase module) from both *S. cerevisiae* and *H. sapiens*. We find that neither holo nor core human TFIIH exhibit processive translocation, consistent with the lack of start-site scanning in humans. Furthermore, in contrast to holo-TFIIH, the *S. cerevisiae* core-TFIIH also lacks processive translocation and its dsDNA-stimulated ATPase activity was reduced ~5-fold to a level comparable to the human complexes, potentially explaining the reported upstream shift in start-site observed in the absence of the *S. cerevisiae* kinase module. These results suggest that neither human nor *S. cerevisiae* core-TFIIH can translocate efficiently, and that the *S. cerevisiae* kinase module functions as a processivity factor to allow for robust transcription start-site scanning.

## Introduction

The general transcription factor TFIIH plays a pivotal role in both RNA polymerase II (Pol II) transcription initiation and DNA repair[1–3]. The core TFIIH complex from *H. sapiens* and other metazoans has seven subunits: XPB, XPD, p62, p52, p44, p34, and p8. The core is functional in nucleotide excision repair (NER) and is dependent on the XPB and XPD nucleic acid motor proteins to unwind DNA around a site of damage for excision and repair[4]. For transcription initiation, core-TFIIH associates with a kinase module comprised of three proteins: cyclin-dependent kinase 7 (CDK7), Cyclin H and MAT1, thus forming holo-TFIIH. In metazoans, the TFIIH kinase module is also called the CDK activating kinase (CAK), because CDK7 activates other cellular CDKs[3,5]. The orthologous 3-subunit TFIIH kinase module in yeast (*S. cerevisiae*) is called TFIIK. Similar to metazoan TFIIH, TFIIK associates with core-TFIIH to form a 10-subunit holo-TFIIH complex. During transcription initiation and promoter escape, CDK7 phosphorylates the C-terminal domain of the Pol II Rpb1 subunit[6,7]. Unlike in DNA repair, XPD only plays a structural role in the formation of the pre-initiation complex (PIC)[8], while XPB activity is required for promoter unwinding[9]. While these molecular activities appear to be generally conserved across Eukaryotes, in yeast exclusively (*S. cerevisiae,* and a host of recently identified related yeasts), TFIIH drives transcription start-site (TSS) scanning and generates a distinct TSS profile genome-wide compared to other Eukaryotes[10–13].

The TSS-scanning activity of the yeast Pol II PIC results in downstream TSS being favored compared to the upstream site of initial DNA unwinding. More specifically, the *S. cerevisiae* PIC utilizes start-sites 40-150 bp downstream of the TATA box whereas metazoans initiate transcription 25-30 bp downstream of the TATA sequence[10,14–16]. Yeast and human Pol II initiation complexes share a high degree of structural conservation with no apparent distinctions that explain differences in TSS-scanning propensity[17,18]. In studies of the *S. cerevisiae* homologue of XPB, Ssl2, we previously showed that in the context of TFIIH, scanning depends on an ATP-dependent double-stranded DNA (dsDNA) translocase activity which leads to a partial DNA opening[19,20]. Furthermore, mutations in the Pol II trigger loop (causing faster or slower transcription kinetics) shift the TSS upstream or downstream, respectively[13,21]. Taken together, these observations suggest that a kinetic competition between Pol II initial transcription and TFIIH dsDNA translocation determines TSS usage[13,20]. Specifically, this model predicts that when TFIIH translocation is faster than initial transcription, downstream scanning will occur. Conversely, if translocation kinetics are relatively slow, initiation at upstream sites will be favored.

In light of this kinetic competition model, we hypothesized that the difference could be explained by distinct XPB/Ssl2 ATPase kinetics or motor properties. To test this hypothesis, we determined the parameters of dsDNA translocation of both *H. sapiens* and *S. cerevisiae* TFIIH complexes in the presence and absence of their respective kinase modules. Consistent with our hypothesis, we find that human TFIIH is a poor dsDNA translocase compared to *S. cerevisiae* TFIIH and that the yeast kinase module specifically stimulates processive translocation.

## Results

### *Hs*TFIIH XPB ATPase is stimulated by double-stranded (ds) DNA

As described above, *S. cerevisiae* TFIIH has been shown to have an ATP-dependent dsDNA translocase activity which is required for DNA unwinding and scanning[19]. In contrast, scanning does not appear to occur in metazoans at TATA-dependent promoters despite structural conservation between *S. cerevisiae* and *H. sapiens* Pol II PICs[17,18]. One possible explanation for this lack of scanning could be a difference in the dsDNA translocase kinetics of TFIIH. To see if the *H. sapiens* TFIIH (*Hs*TFIIH) dsDNA translocase kinetics differs from *S. cerevisiae* TFIIH (*Sc*TFIIH), we quantitated the steady-state ATPase kinetics as a function of DNA length as done previously with the *S. cerevisiae* enzyme[19].

TFIIH ATPase activity was measured using a continuous fluorescence assay in plate reader format in which ATP turnover is coupled to NADH reduction through the action of pyruvate kinase and lactate dehydrogenase (Fig. 1A **and Methods**). For every equivalent of ADP generated, an equivalent of NADH is oxidized to NAD, resulting in a decrease in fluorescence[22,23]. The continuous assay is advantageous, yielding more data points in a lower reaction volume and adapting the assay to a plate format allows for multiple experiments to be performed in parallel. Core- or holo-*Hs*TFIIH was preincubated in the absence of DNA or the presence of either plasmid DNA (5500 bp) or linear dsDNA in *Hs*-ATPase buffer at 30 °C for 10 min. ATP and phosphoenol pyruvate (PEP) were added to initiate the reaction. No ATPase activity was detectable in the absence of DNA, but in the presence of DNA, the fluorescence signal decreased linearly over time (Fig. 1B). This fluorescence signal was converted to moles of ADP using a standard curve (collected during each experiment; Fig. 1C) and fit to a line to measure steady-state ADP production (Fig. 1D). The plasmid-concentration dependence of the core-*Hs*TFIIH steady-state rate increased hyperbolically, displaying Michaelis-Menten kinetics with K_m_ = 12 ± 3 μM-bp and V_max_ = 0.8 ± 0.1 ATP/sec/TFIIH (Fig. 1E). Holo-*Hs*TFIIH produced similar linear ATPase time courses at saturating plasmid concentration (Fig. 1D); however, the V_max_ was approximately 2-fold lower (V_max_ = 0.5 ± 0.02 ATP/sec/TFIIH).

**Figure 1.**
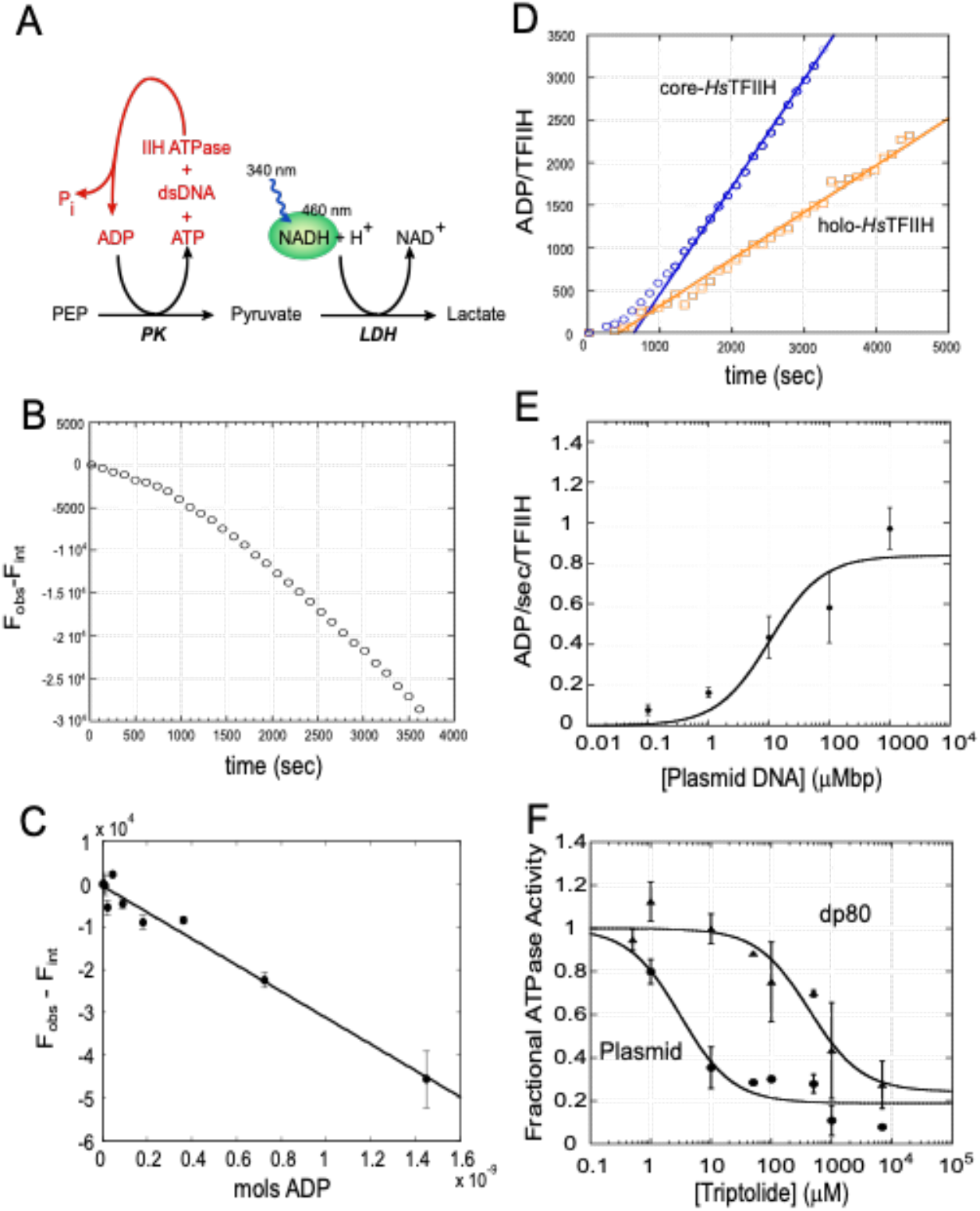
Human TFIIH core ATPase is stimulated by dsDNA and is XBP-dependent. **(A)** Diagram of the continuous, fluorescent, enzyme-coupled ATPase assay used to measure TFIIH steady-state ATPase activity. TFIIH ATP hydrolysis was monitored by following the oxidation of NADH to NAD through the regeneration of ADP to ATP through the glycolytic enzymes pyruvate kinase and lactate dehydrogenase. NADH fluorescence at 460 nm is quenched upon NADH oxidation to NAD. **(B)** Fluorescence time course of core-*Hs*TFIIH ATPase activity in the presence of plasmid DNA after subtracting control time course of core-*Hs*TFIIH in the absence of DNA. **(C)** ADP standard curve for converting observed NADH fluorescence to moles of ADP. **(D)** Representative ATPase time courses for core- and holo-*Hs*TFIIH. The core-*Hs*TFIIH, time course is the same time course in (B) converted to ADP/*Hs*TFIIH using the standard curve in (C) and the moles of *Hs*TFIIH determined by SDS-PAGE as described in Materials and Methods. The slope of a linear fit is the observed rate of ATP hydrolysis. **(E)** The rate of core-*Hs*TFIIH ATP hydrolysis as a function of plasmid DNA concentration. The trends are fit with the Michaelis-Menten equation (solid line), see Materials and Methods. **(F)** Triptolide inhibition curves for core-*Hs*TFIIH ATPase in the presence of plasmid (circles) and linear dsDNA (80 bp, triangles).

To distinguish between the ATPase activities of XPD and XPB, we used the XPB-specific inhibitor triptolide[24] **(Supplemental Fig. 1A)**. Core-*Hs*TFIIH was incubated with 100 μM-bp plasmid DNA in the absence or presence of increasing amounts of triptolide in Hs-ATPase buffer at 30 °C for 20 minutes prior to reaction. As judged by fitting the data to an inhibitory binding curve, 81 ± 4% of the observed ATPase activity was inhibited by triptolide, indicating the plasmid DNA-stimulated ATPase activity is XPB-dependent (Fig. 1F). An 80 bp linear dsDNA was also tested since the presence of a free DNA end, although blunt, could stimulate loading of XPD, resulting in ATP hydrolysis[25]. As with the plasmid template, triptolide inhibited the majority (80 ± 10%) of the observed ATPase activity on the 80 bp dsDNA (Fig. 1F) although the inhibition curve is shifted to higher triptolide concentrations. As TFIIH was incubated with triptolide in the presence of DNA, this suggested that DNA may competitively inhibit triptolide binding to core-*Hs*TFIIH and that core-*Hs*TFIIH has a higher affinity for the 80 bp linear DNA than the plasmid DNA.

Structures of XPB in the contexts of core-TFIIH and the PIC provide insight into how triptolide disrupts its ATPase activity and how DNA binding may compete with triptolide binding. Mass spectroscopic analysis of triptolide inactivated-*Hs*TFIIH showed cysteine 342 of XPB to be covalently modified[26]. Inspection of the apo core-*Hs*TFIIH structure[27] reveals cysteine 342 to be exposed to solvent and positioned at the back of a pocket surrounded by residues that surround and form the ATP binding site in nucleic-acid-stimulated ATPases[28]. However, in the DNA-bound XPB sub-structure from the human PIC, cysteine 342 becomes less solvent exposed due the presence of the bound DNA[17] **(Supplemental Fig. 1B)**. This can be seen directly by aligning the apo and DNA bound XPB structures and is consistent with our data which suggests that bound DNA limits triptolide access to cysteine 342 **(Supplemental Fig. 1C)**. Speculatively, in turn, a triptolide adduct at cysteine 342 would be predicted to disrupt both ATP- and DNA-binding by XPB.

### Both core- and holo-*Hs*TFIIH are poor dsDNA translocases

The dsDNA-length dependence of the steady-state ATPase kinetic parameters, K_m_ and V_max_, can provide insight into the translocation mechanism of nucleic acid motors[19,29,30]. In particular, the probability of the motor to take a step along the DNA rather than dissociate, quantitated by the processivity (*P*), can be determined from the DNA length dependence of V_max_[30]. For processive motors, V_max_ will increase as the DNA length increases past the average length translocated per binding event, and plateau at significantly longer lengths. In contrast, a DNA-length independent V_max_ provides evidence of either no processivity or a processivity significantly less than the lowest DNA length tested. We measured the steady-state ATPase rate for core-*Hs*TFIIH, titrating the concentration of a series of linear dsDNA substrates with different lengths (20, 30, 60, 80 and 100 bp). These data were compared with plasmid DNA, representing an infinitely long template. The steady-state rate increased hyperbolically with DNA concentration for each DNA length tested (Fig. 2A). Each titration was fit to the Michaelis-Menten equation to determine V_max_ and K_m_. In contrast to a length-dependent V_max_ previously observed using holo-*Sc*TFIIH[19], V_max_ was DNA length-independent with core-*Hs*TFIIH (Fig. 2B, **blue**). The K_m_ was similar for lengths 20, 30, 60, 80, and 100 bp, but plasmid DNA K_m_ was 2-fold higher, suggesting a lower affinity toward plasmid DNA (Fig. 2C).

**Figure 2.**
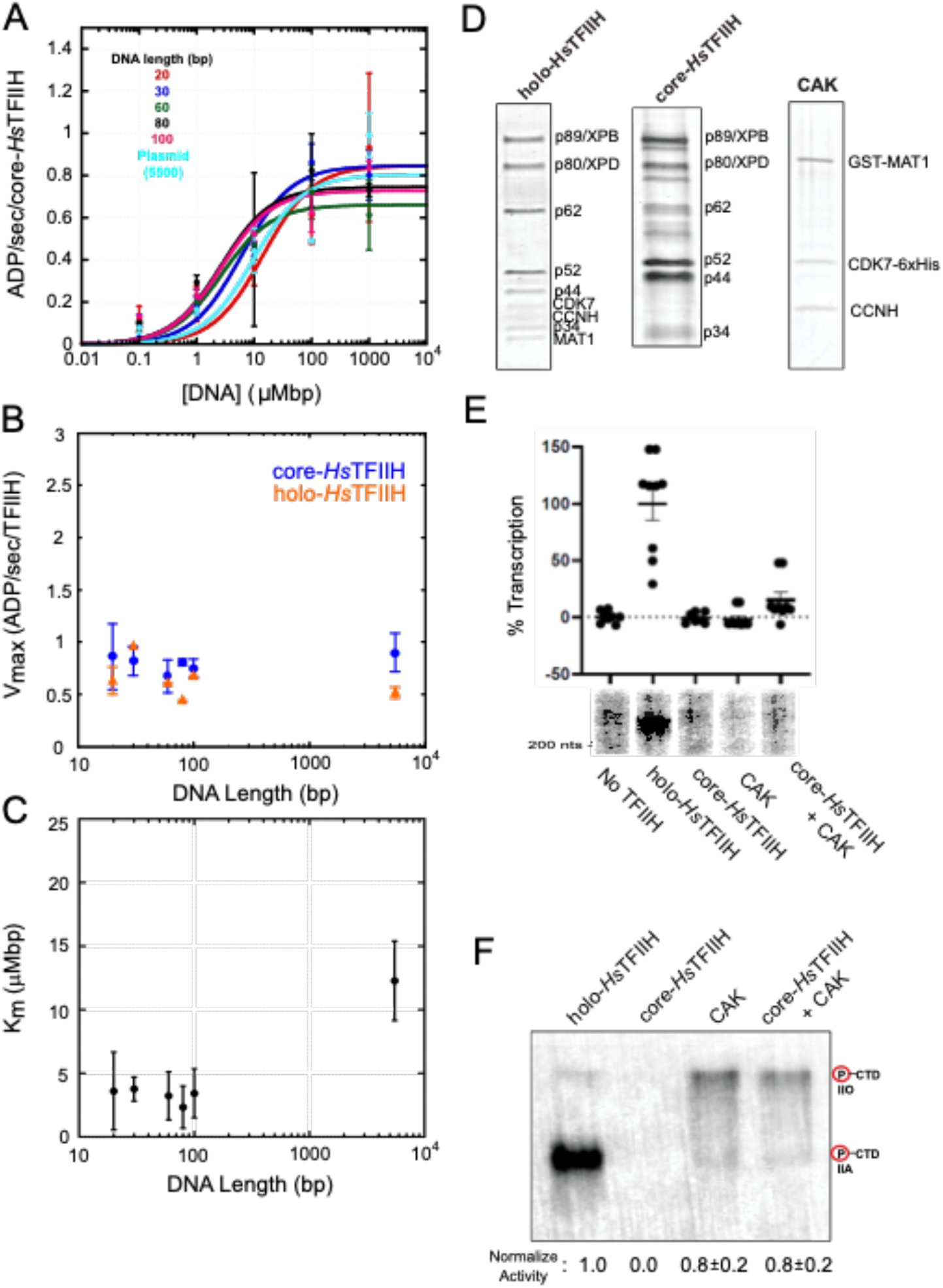
Core- and holo-*Hs*TFIIH are poor dsDNA translocases. **(A)** DNA concentration dependence of core-*Hs*TFIIH steady-state ATPase rate for different linear dsDNA lengths (lengths: 20, 30, 60, 80 and 100 bp) and plasmid DNA. Data was fit with the Michaelis-Menten equation (solid lines) to determine K_m_ and V_max_ for each DNA length. **(B)** Core-*Hs*TFIIH and holo ATPase V_max_ as a function of DNA length. A V_max_ independent of DNA length suggests very low to no processivity for dsDNA translocase. **(C)** Core-*Hs*TFIIH ATPase K_m_ for DNA as a function of DNA length. **(D)** SDS-PAGE sypro ruby stain of purified core and holo-*Hs*TFIIH complexes and commercially purchased CAK. **(E)** *In vitro* transcription of *Hs*TFIIH complexes with or without added CAK. (**F**.) *In vitro* kinase assay of *Hs*TFIIH complexes with or without added CAK. Both a hyperphosphorylated (IIO) and hypo-phosphorylated (IIA) CTD is observed.

We wondered if the lack of length-dependence could be explained by the lack of the kinase module, which has been suggested to play a role in yeast TSS-scanning. To test this idea, we determined the V_max_ for holo-*Hs*TFIIH at saturating DNA concentration (1000 μM-bp) for each DNA length. V_max_ for holo-*Hs*TFIIH was slightly lower than core-*Hs*TFIIH and still DNA length independent (Fig. 2B, **orange)**. The lack of DNA-length dependence for both core- and holo-*Hs*TFIIH, suggests that, in contrast to holo-*Sc*TFIIH, human TFIIH complexes are not processive dsDNA translocases. Importantly, all subunits of holo-*Hs*TFIIH are present in the preparation (Fig. 2D) and the complex was active in both transcription and kinase assays (Fig. 2E **and** F), as expected. In contrast, despite having the expected subunits and displaying ATPase activity, Core-*Hs*TFIIH was not active in transcription assays (Fig. 2E). Furthermore, when commercially available purified CAK (Millipore, cat: 14-476M, CDK7-6xHis/CCNH/GST-MAT1) was added to reactions with core-*Hs*TFIIH the “core+CAK” activity did not match that of purified intact holo-TFIIH (Fig 2E, F), suggesting that it was not incorporated into a *bona fide* holo-complex.

### Removal of the *S. cerevisiae* kinase module reduces both ATPase rate and dsDNA translocation processivity

As described above, the kinase module had little effect on *Hs*TFIIH dsDNA translocase activity. However, the removal of the *S. cerevisiae* kinase module has been shown result in upstream shifts in TSS[31]. In fact, this shift leads to the usage of a start-site position similar to that found in metazoans. While the shift is not absolute, the result suggests a significant reduction in the prevalence of TSS scanning. Furthermore, adding back an enzymatically dead kinase module restores usage of downstream start-sites, suggesting that the module plays a structural role, as opposed to a catalytic one, in facilitating TSS scanning[31]. We hypothesized that if scanning activity is due to processive TFIIH translocation as in the kinetic competition model, removal of the kinase module should also reduce (or eliminate) *Sc*TFIIH dsDNA translocase activity.

To test this, we affinity purified core-*Sc*TFIIH (Fig. 3A) and measured the steady-state dsDNA-stimulated ATPase activity of core- and holo-*Sc*TFIIH at saturating DNA concentrations on DNA templates of varying lengths as described above. As the Tfb3/MAT1 subunit of the kinase module anchors the kinase module to core-TFIIH[32], core-ScTFIIH was purified from a strain containing Tfb3 with an auxin degron tag (see Methods). Both core and holo complexes also contained an ATPase-dead Rad3 subunit (homologous to XPD) to ensure that any ATPase activity was from Ssl2 (homologous to XPB). Furthermore, both holo- and core-*Sc*TFIIH were active in transcription where PIC formed with core-*Sc*TFIIH was able to initiate transcription upstream at or near the metazoan start site with less initiation at downstream sites as previously observed (Fig. 3B)[31]. As expected from our previous work, the ATPase time courses for holo-*Sc*TFIIH increase linearly (Fig. 4A) and V_max_ increases with DNA length, plateauing at lengths greater than 100 bp (Fig. 4B, **orange)** [19]. A non-linear least-squares analysis of the DNA-length dependence of V_max_ yielded a processivity of 0.91 ± 0.01 equivalent to on average 11 bps translocated per binding event.

**Figure 3.**
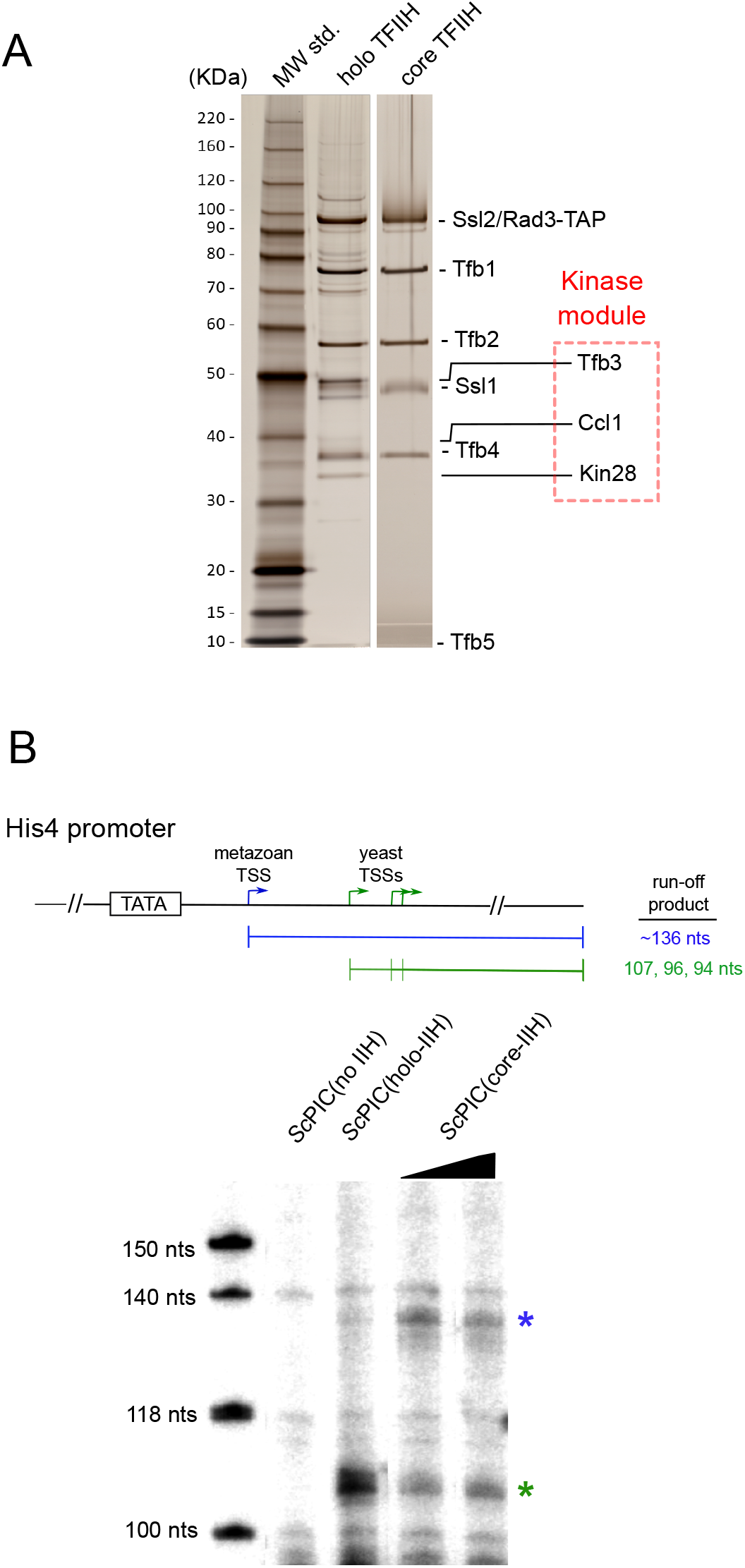
Purified core-*Sc*TFIIH displays impaired TSS-scanning. **(A)** SDS-PAGE silver stain of purified holo- and core-*Sc*TFIIH. **(B)** *In vitro* run-off transcription with holo- and core-*Sc*TFIIH on the HIS4 promoter template. Asterisks denote RNA products initiated from the metazoan (blue) and yeast primary endogenous (green) transcription start-sites. Secondary TSSs from the yeast reaction are obscured by an artifact band also observed in the no TFIIH control lane.

**Figure 4.**
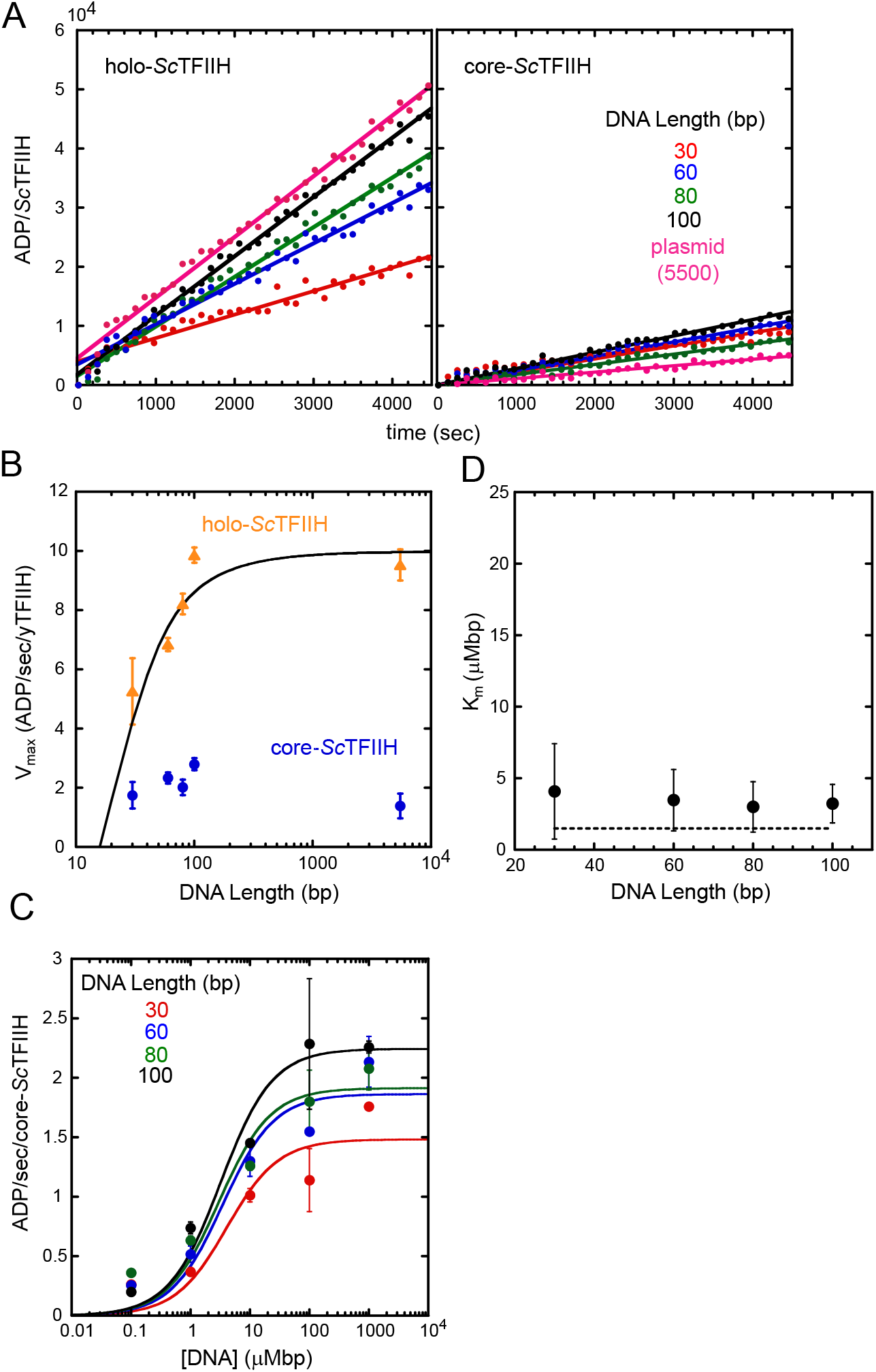
*Sc*TFIIH kinase module promotes dsDNA translocation processivity and enhances ATPase activity. **(A)** Steady-state ATPase time courses for holo- and core-*Sc*TFIIH at saturating DNA (1000 μM-bp) for linear DNA (lengths: 30, 60, 80, and 100 bp) and plasmid DNA. **(B)** Holo- and core-*Sc*TFIIH ATPase V_max_ as a function of DNA length. Holo-*Sc*TFIIH V_max_ as a function of DNA length was fit with equation (1) to determine the translocation processivity. Core-*Sc*TFIIH core V_max_ is independent of DNA length, indicating the translocase has very low to no processivity. **(C)** DNA concentration dependence of core-*Sc*TFIIH steady-state ATPase. Data was fit with the Michaelis-Menten equation (solid lines) to determine V_max_ and K_m_. **(D)** Core-*Sc*TFIIH K_m_ as a function of DNA length. Dashed line is the average K_m_ (1.5 ± 0.3 μM-bp) previously observed for holo-*Sc*TFIIH[19] under the same solution conditions.

In contrast, core-*Sc*TFIIH time courses increase linearly at a slower rate (Fig. 4A) and the measured V_max_ values do not show DNA-length dependence (Fig. 4B, **blue**). This result is consistent with core-*Sc*TFIIH lacking observable dsDNA translocase activity similar to the behavior of the human complexes. We also measured the core-*Sc*TFIIH DNA concentration dependence of the ATPase activity to confirm that we were measuring V_max_ in the presence of saturating DNA and to eliminate alternative models of translocation that predict a length-dependence in K_m_ instead of V_max_[30]. Each DNA length tested displayed a hyperbolic dependence on DNA concentration which was fit to the Michaelis-Menten equation to determine K_m_ (Fig. 4C). The core-*Sc*TFIIH K_m_ was similar for each length (Fig. 4D) and higher than holo-*Sc*TFIIH (K_m_ = 1.5 ± 0.3 μM-bp) determined previously (Fig. 4D dashed line)[19], suggesting the core-*Sc*TFIIH has a lower affinity for DNA than holo-*Sc*TFIIH. Interestingly, the core-*Sc*TFIIH V_max_ is 5-fold lower than the holo-*Sc*TFIIH V_max_ at long DNA lengths revealing that the kinase module not only promotes dsDNA translocation processivity but also appears to enhance the absolute dsDNA stimulated ATPase rate. Taken together, these results further support a mechanism in which TFIIH dsDNA translocation is the underlying activity that produces TSS-scanning in *S. cerevisiae*.

## Discussion

The general transcription factor TFIIH plays a pivotal role in Pol II transcription initiation, facilitating promoter DNA opening by translocating downstream DNA into Pol II. As TBP binds upstream DNA, this translocation generates torsional strain which then promotes DNA melting around the TSS. This process is conserved among Eukaryotes and allows ssDNA to enter the Pol II active site. However, in some yeast species, TFIIH also facilitates TSS-scanning in which, after initial DNA unwinding, the PIC scans downstream DNA and utilizes TSS located 40 – 120 bp downstream of TATA. This scanning mechanism requires Ssl2/XPB, the ATP-dependent dsDNA translocase of TFIIH. The current model for scanning proposes a kinetic competition between Ssl2/XPB dsDNA translocation and Pol II initial transcription. This model is supported by the requirement of TFIIH dsDNA translocation and the observation that loss-of-function mutations in *S. cerevisiae* Pol II (i.e. slower polymerase) shift the TSS downstream, whereas gain-of-function mutations (i.e. faster polymerase) shift the TSS upstream[13]. Interestingly, metazoan Pol II PICs do not appear to scan on TATA containing promoters and instead utilize TSS that are 25-30 bp downstream of the TATA sequence[10,14,16]. In the context of the kinetic competition model, a lack of scanning would be predicted in the case of a shift in the relative kinetics between the Pol II and Ssl2/XPB motors due to faster Pol II initial transcription and/or slower Ssl2/XPB dsDNA translocation kinetics. Therefore, we undertook the work described here to test the hypothesis that metazoan TFIIH is a slower and/or less processive dsDNA translocase compared with *S. cerevisiae* TFIIH.

We previously showed that *S. cerevisiae* holo-TFIIH processively translocates along dsDNA using a discontinuous ATPase assay[19]. We repeated those experiments using the continuous fluorescent ATPase assay used to measure core- and holo-*Hs*TFIIH in this study and obtained results consistent with our previous study. Analysis of the DNA-length dependence of the ATPase V_max_ yielded a translocation processivity of 0.91 (11 bp per binding event on average), indicating holo-*Sc*TFIIH translocates about one turn of dsDNA, on average, before dissociation from DNA (Fig. 4B). In contrast, neither core- nor holo-*Hs*TFIIH showed any length-dependence of V_max_ for the DNA lengths tested. This result indicates that their processivity, if any, cannot be detected by this assay (Fig. 2B). However, we can place an upper limit on the processivity of 2 bp by comparing the data to V_max_ values calculated from equation (1) for different DNA lengths and varying processivities. This analysis shows that processivities of 1 and 2 bp are within the error of the observed V_max_ values **(Supplemental Fig. 2)**. Crucially, the affinity purified holo-*Hs*TFIIH was active in transcription and kinase assays *in vitro,* suggesting the lack of processivity is a *bona fide* feature of the complex. Furthermore, the observed ATPase activity in *Hs*TFIIH was inhibited by triptolide, establishing that XPB, and not XPD, was the motor stimulated in our assays. These observations reveal that holo-*Hs*TFIIH is a poor DNA translocase compared to holo-*Sc*TFIIH. These data are consistent with the kinetic competition model, in which the rate of Pol II initial transcription vs. TFIIH-dependent DNA translocation determine TSS usage. Indeed, kinetic simulations that vary translocase processivity relative to initial transcription processivity (Fig. 5A) result in TSS distributions that expand downstream with increasing translocase processivity (Fig. 5B).

**Figure 5.**
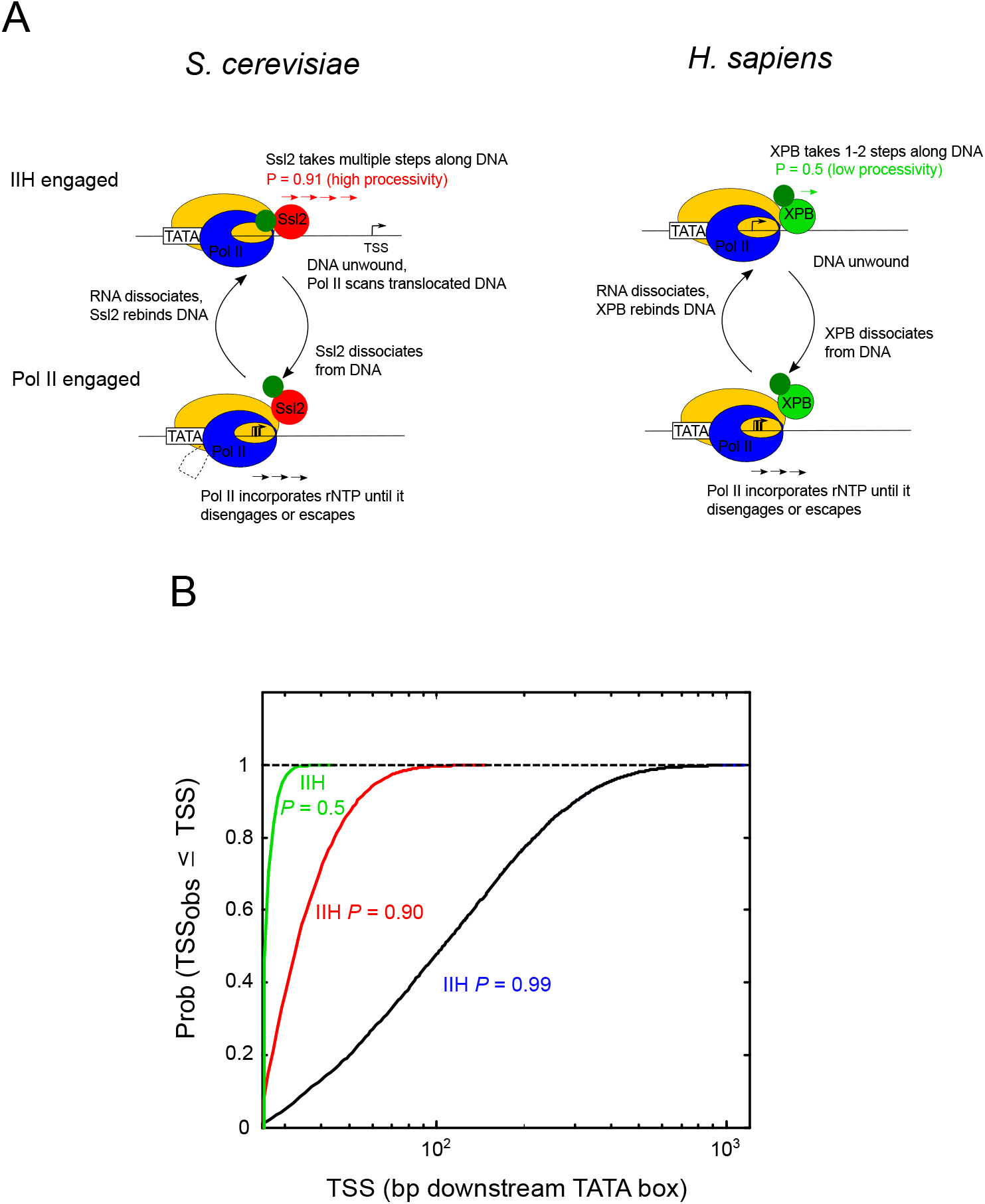
TFIIH dsDNA translocase processivity limits TSS-scanning. **(A)** A kinetic competition model for TSS-scanning comparing the *S. cerevisiae* and *H. sapiens* systems. When TFIIH is engaged with the DNA and translocating, Pol II is prohibited from initiating transcription. When TFIIH dissociates from the DNA Pol II initiates transcription. If a 10-mer RNA is formed a transcription start site is scored. See Material and Methods for details on the model and simulations. Ssl2 is more processive and takes more steps (red arrows) downstream than XPB (green arrow) per binding event. The kinase module (dark green) is depicted closer to the DNA and Ssl2 during translocation to indicate a direct effect on Ssl2 translocation. The dashed line indicates DNA that must be extruded between TBP and the polymerase during scanning. In contrast, the kinase module has no effect on XPB translocation. **(B)** Cumulative distribution functions of transcription start sites from simulations of the kinetic competition model for TSS-scanning with different processivities for TFIIH dsDNA translocation (P = 0.5 (~2 bp/binding), 0.9 (~10 bp/binding), 0.99 (~100 bp/binding)) and a constant Pol II processivity (0.98).

Intriguingly, PICs formed with core-*Sc*TFIIH have a significantly reduced ability to scan downstream to endogenous TSS and instead initiate upstream near the TSS observed in metazoans[31]. This result suggested that removal of the kinase module may reduce the processivity and/or translocase activity of *Sc*TFIIH. Our assays bore this out: core-*Sc*TFIIH complexes displayed a DNA-length independent V_max_ and a 5-fold lower rate than holo-*Sc*TFIIH at long DNA lengths (Fig. 4B). In addition, PICs formed with core-*Sc*TFIIH in our assay displayed upstream shifted TSS consistent with previous observations (Fig. 3B)[31]. These observations support the kinetic competition model of TSS scanning.

The TFIIH kinase module itself is not well-resolved in the current PIC and TFIIH structural models[18,27,33,34]. However, a number of chemical cross-links have been observed between Tfb3/MAT1, the main component of the kinase module that connects it to core-*Sc*TFIIH, and Ssl2/XBP[32]. These cross-links are located away from the Ssl2 DNA binding cleft and similar cross-links were observed in *Hs*TFIIH. Thus, while a Tfb3/MAT1 interaction with the Ssl2/XPB N-terminal region could potentially stimulate *S. cerevisiae* Ssl2 ATPase and translocase processivity through an allosteric mechanism, the presence of these contacts in both species argues that these contacts do not play direct roles in TSS scanning.

Based on Endonuclease III digestions in the presence and absence of the kinase module, it has been suggested that the kinase module affects the PIC footprint on the promoter DNA[31]. Whether this is via direct DNA binding or indirect changes in the footprints of core-TFIIH or the polymerase is not known. However, if this conformational change prevents the *S. cerevisiae* PIC from initiating at the upstream metazoan site and/or stimulates the processivity of Ssl2, it may partially underlie the observed shift in TSS.

The effect of the *S. cerevisiae* kinase module on dsDNA translocation described here adds to observations in other contexts that the nucleic acid motors within core-TFIIH can be activated through factor-specific interactions. More specifically, core-*Hs*TFIIH DNA translocation activity was shown to be enhanced by the NER factor XPA although XPA-dependent activation was not unambiguously assigned to either XPB or XPD[35]. Relatedly, structures of core-*Hs*TFIIH in the presence and absence of a partial kinase module (MAT1) revealed that core subunits interact with XPD and may block its DNA binding sites, suggesting that conformational rearrangements stimulated by protein-protein interactions, are needed to activate XPD[27,34].

While the removal of the kinase domain from *Sc*TFIIH results in loss of a processivity signal in our assay (Fig. 4B), the core-*Sc*TFIIH still retains some ability to scan to the endogenous sites as judged from the *in vitro* transcription reaction itself (Fig. 3B). This suggests that the activities of TFIIH may be modulated by its incorporation into the PIC. One consistent interpretation of these data is that the processivities of both core- and holo-*Sc*TFIIH are stimulated by protein-protein contacts with PIC general transcription factors. Future studies will be needed to determine how interactions between XPB/Ssl2, the kinase module, and other general transcription factors modulate the molecular activities we have observed here.

## Material and Methods

### Protein Purification

Core-*Hs*TFIIH complex (XPB-preScission-GFP, XPD, p62, p52, p44, p34, and p8) was cloned into MacroBac vector 438a[36]. The protein was expressed in Sf9 cells supplemented with 1 mM L-cysteine and 0.1 mM ferric ammonium citrate. Based on the XPB fusion with GFP, TFIIH was purified anaerobically using GFP-nanobody binder[37] covalently linked to agarose-beads (NHS agarose, Pierce/Thermo); eluted with preScission protease; and purified using Superpose 6 (10/300) in 25 mM HEPES, pH 7.8, 150 mM NaCl, 50 mM KCl, 3% glycerol, and 3 mM β-mercaptoethanol. The elution with TFIIH core complex (confirmed by SDS-PAGE gel and 420 nm absorption peak) was stored at –80 °C until use. The holo-*Hs*TFIIH purification was completed as described[38].

Core-*Sc*TFIIH was purified from strain SHY1290 (genotype: *mat a ade2::hisG his3 Δ200 leu2Δ0 lys2Δ0 met15Δ0 trp1Δ63 ura3Δ0 rad3Δ*::KanMx *tfb6Δ*::HPH p*GPD1*-OSTIR::*HIS3 TFB3*-3xV5-IAA7::*URA3/*pJF82 (ars cen *LEU2 rad3* (E236Q) – (HA)1-Tap tag). This strain contains a Tap-tagged ATPase dead Rad3/XPD (E236Q); a deletion of the *TFB6* gene to facilitate complex purification; and Tfb3 with an auxin degron tag to facilitate elimination of the kinase module. Cells (6 liters) were grown at 30 °C in YPD + adenine media (3% final Glucose) to Abs600 = 3-4. 3-indoleacetic acid (IAA) in DMSO was added to 0.5 mM final for 30 min to deplete Tfb3 protein (the protein was undetectable by Western blot after this treatment). Core-*Sc*TFIIH and holo-*Sc*TFIIH with an ATPase dead Rad3/XPD were purified by Tap-tag as previously described[19]. Core-*Sc*TFIIH was additionally purified by chromatography on a 0.28 ml Source 15Q column equilibrated in 20 mM HEPES, pH 7.6, 10% (v/v) glycerol, 200 mM KOAc, 1 mM EDTA, and 1 mM DTT. Protein was eluted over 30 column volumes with a KOAc gradient from 200 mM to 1.2 M. Fractions with core-*Sc*TFIIH were pooled and loaded on a 0.1 ml calmodulin affinity column, as previously described[19], to remove trace amounts of Ccl1 and Kin28 of the kinase module. Fractions were assayed by silver stain and those containing core-*Sc*TFIIH were pooled, aliquoted, and stored at –80 °C. The concentration of the TFIIH complexes were determined by comparing the purified protein with a titrated BSA standard on SDS-PAGE gel (*Hs*TFIIH) or by quantitative Western (*Sc*TFIIH).

The *S. cerevisiae* Pol II RNA polymerase and the other general transcription factors (TBP, TFIIB, TFIIF, and TFIIE) were purified as previously described[39]. The human factors (Pol II, TBP, TFIIB, TFIIF, TFIIE, and TFIIH) were purified as described[38].

### DNA preparation

DNA oligos of a random sequence and corresponding complementary oligos were purchased from IDT (Integrated DNA Technologies, Inc., Coralville, Iowa, USA) and resuspended in 10 mM Tris-HCl, pH 8.0. For a given DNA length oligos were annealed in a 1:1 molar ratio in Annealing buffer (10 mM Tris-HCl, pH 8.0, 50 mM NaCl). The DNA solution was heated at 95°C for 5 min then slowly cooled (~4-5 hrs) to room temperature (~23 °C) in an insulated heating block. Annealed DNAs were verified by native-PAGE following standard protocols. The annealed DNA concentrations were determined from their absorbance at 260 nm using the calculated molar extinction coefficient for the duplex DNA based on the DNA sequence (nearest neighbor ref). A midi prep scale of plasmid DNA (pGO10GG, 5500 bp, sequence available upon request) was purified following manufacture protocol (Qiagen). The promoter template for in vitro transcription with the human Pol II system was prepared from the native human HSPA1B gene. The HSPA1B gene was initially amplified from HeLa genomic DNA by PCR (forward primer: CTCCTT CCCATT AAGACG GAAAAA ACATCC GGGAGA GCCGGT CCG; reverse primer: ACCTTG CCGTGT TGGAAC ACCCCC ACGCAG GAGTAG GTGGTG CCCAGGTC) and cloned into a TOPO vector. The −500 to +216 region corresponding to the HSPA1B promoter was amplified from the plasmid by PCR with Phusion polymerase (Thermo-Fisher #F530S) and purified via E.Z.N.A Gel Extraction Kit (Omega BioTek #D2500). The DNA was ethanol precipitated, resuspended to 100 nM in milliQ water, and stored at −80 °C in 10 μL single-use aliquots. The promoter template for in vitro transcription with the *S. cerevisiae* Pol II system was prepared from the native His4 gene. A 236 bp DNA region, spanning −74 to +107, was PCR amplified from plasmid (pSH515) containing the native His4 gene promoter and purified by PCR cleanup kit (Qiagen). DNA was eluted from the spin column with 10 mM Tris-HCl, pH 8.0 and stored at −20 °C.

### Fluorescent coupled ATPase assay

Steady-state ATP hydrolysis was measured by coupling the production of ADP to the reduction of NADH via the glycolytic enzymes pyruvate kinase and lactate dehydrogenase as depicted in Figure 1A[22]. ATPase reactions with *Hs*TFIIH were carried out in hTRxn buffer (10 mM Tris-HCl pH 7.9, 10 mM HEPES pH 7.9, 50 mM KCl, 4 mM MgCl_2_, 10% (v/v) glycerol, 0.1 mg/ml BSA, and 1 mM DTT) at 30°C. ATPase reactions with *Sc*TFIIH were carried out in yTRxn buffer (10 mM HEPES pH 7.7, 100 mM KGlu, 10 mM MgOAc_2_, 3.5% (v/v) glycerol, 0.05 mg/ml BSA, and 1 mM DTT) at 25°C. All reagents were diluted and or prepared fresh in the appropriate TRxn buffer the day of the experiment and kept on ice. All reactions were prepared in a total volume of 10 uL the reported concentrations are the final concentrations after mixing. In a 8.75 μL volume, 0.133 mg/ml NADH (β-nicotinamide adenine dinucleotide reduced, Sigma, cat: N4505), 5.6 U/ml pyruvate kinase/ 7.8 U/ml lactate dehydrogenase (Sigma, cat: P0294), TFIIH (1.2 nM yeast variants or 6 nM human variants), and DNA were mixed and incubated on ice for 30 min. The reactions were then transferred to a black, 384 low volume, round bottom well plate (Corning, cat: 4514); covered with optical film to prevent evaporation (Applied Biosystems, cat: 4360954); and placed in the Synergy2 fluorescent plate reader (Bio Tek, Winooski, VT) and incubated for 10 min at the reaction temperature. After an initial scan of the plate (excitation: 340 nm , emission: 460 nm), the film was temporarily removed and the reaction initiated by adding 0.31 mg/ml PEP (phosphoenolpyruvic acid, Sigma, cat: 860077) and 0.5 mM ATP in a 1.25 μL volume using a multichannel pipet. The plate was then rapidly (~5-10 sec) resealed and placed back in the plate reader. Every two minutes the plate was scanned over a period of 75 mins. Triptolide inhibition of the ATPase activity was assayed by incubating TFIIH:DNA with varying amounts of triptolide (Sigma, cat: T3652) at 30 °C for 30 min before initiating the ATPase reaction with PEP and ATP. Triptolide was dissolved in DMSO and stored at −20 °C. The fluorescent signal was calibrated for each plate of reactions by setting up a series of reactions with known amounts of ADP without TFIIH and DNA. All reactions were prepared in triplicate.

### In vitro Run-off transcription assay

Human pre-initiation complexes were assembled on an activated HSPA1B promoter template (5nM in 10 μL) by incubating 400 nM HSF1 with promoter template for 30 minutes at 30°C in transcription buffer (20 mM HEPES pH 7.6, 1 mM DTT, 8 mM MgCl2) and enough DB(100) buffer (10% glycerol, 10 mM Tris pH 7.9, 180 mM KCl, 1 mM DTT) to ensure the final reaction volume is 20 μL. Pre-initiation complex (PIC) mix (TFIIB, TBP, TFIIF, and Pol II) is then added to the activated template with the same amount of TFIIH variants added separately and incubated for 15 minutes at 30 °C to permit PIC assembly. S. cerevisiae pre-initiation complexes were assembled on the His4 promoter template (10 nM) in yTRxn buffer in a final volume of 20 μL at 25 °C. PIC components (Pol II, TBP, TFIIB, TFIIF and TFIIE) along with either holo- or core-*Sc*TFIIH, at equivalent concentrations, are added to the promoter DNA and incubated for 30 min to permit PIC assembly. For both the human and yeast systems optimal amounts of each factor are added to achieve maximal transcription activity.

Transcription is initiated by the addition of 2 μL of NTP mix (25 mM A/G/UTP, 5 mM CTP, 0.1 mCi α^32^P-CTP, brought to 40uL with DB(100) or 20 μL yTRxn buffer and allowed to proceed for 30 minutes at either 30 °C for the human or 25 °C for the yeast system. Reactions are stopped with the addition of 150 μL of STOP buffer (20 mM EDTA, 200 mM NaCl, 1% SDS) and 350uL of cold 100% ethanol. RNA precipitates at −20 °C overnight. Precipitated RNA is resuspended in formamide loading buffer, boiled, and run on a 40 cm 7% acrylamide sequencing gel. RNA was quantitated through pixel-density analysis of the ^32^P signal in the sequencing gel using ImageJ. Bands corresponding to 216 nt were regarded as products of successful human Pol II transcription and selected for quantitation with background correction. Data from ImageJ was analyzed and graphed with GraphPad Prism8 software. For the yeast system bands corresponding to 138 nt are products initiated from the metazoan start-site while bands corresponding to 107, 96, and 94 nt are products initiated from the native downstream start-sites.

### In vitro kinase assay

The kinase assays evaluating CDK7 activity with the *Hs*TFIIH variants were performed with equivalent concentrations of holo-TFIIH, core-TFIIH, and the CAK with the Pol II CTD as the substrate and 100 μM ATP and ^32^P-γ ATP for 30 minutes at 30°C. These conditions were chosen to remain consistent with the *in vitro* transcription assays. The CAK complex was purchased from Millipore (CDK7-6xHis/CCNH/GST-MAT1; CAK complex, 250 μg cat: 14-476M) and the Pol II GST-CTD was expressed in BL21 cells and purified as described[40]. Kinase reactions were quenched with 4X Laemmli buffer, boiled for 5 minutes, and run on a 4-20% gradient protein gel (BioRad 4–20% Mini-PROTEAN® TGX™ Gel, 15 well, 15 μl cat#456-1096). Gels were dried and exposed to evaluate autorad signal. ImageJ software was used to measure the autorad signal. The measured kinase activity was normalized to TFIIH holo and the standard deviation was reported.

### Kinetic analysis and modeling

The initial ATPase steady-state rate was determined from the slope of the linear portion of the ATPase time courses. The ATPase steady-state kinetic parameters V_max_ and K_m_ for a given DNA substrate was determined by fitting the initial ATPase steady-state rate as a function of DNA concentration to the Michaelis-Menten equation. The steady-state ATPase rate for a translocating motor along a lattice will display Michaelis-Menten kinetics where the V_max_ will change as a function of lattice length[30]. The expression for the DNA length dependent change in V_max_ is given in Equation (1) and was previously used to determine the translocation processivity for holo-*Sc*TFIIH[19].

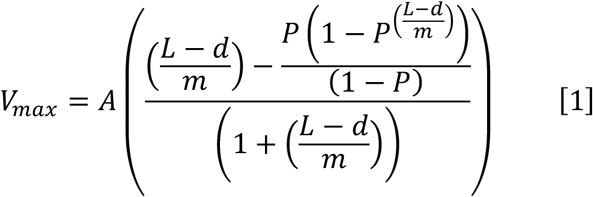

In Equation (1), P is the processivity, L, the DNA length, d, the translocase contact size, and m, the translocase step size per cycle. The parameter A is a product combination of microscopic parameters for the translocation model and the contact size and step size were constrained to 16 bp and 1 bp, respectively as done previously[19] while the processivity was allowed to float in a non-linear least-squares analysis of the DNA length dependence of the V_max_ using Conlin[41].

Sequence independent distributions of transcription start-sites were simulated using the continuous scanning model depicted in Figure 5A. When TFIIH is engaged with the DNA and translocating, Pol II is prevented from initiating transcription. When TFIIH dissociates, Pol II initiates transcription. If it transcribes a 10-mer before dissociating a transcription start-site is scored and the system resets; otherwise, the short RNA is released and TFIIH rebinds and continues translocation along the DNA. Gillespie simulations were performed in Matlab until 10,000 transcription start-sites were scored for different TFIIH processivities (P = 0.50, 0.90, 0.99, k_dissociate_ = 10 s^−1^, 1 s^−1^, 0.1 s^−1^ respectively and k_translocate_ = 10 s^−1^ with a constant Pol II processivity (P = 0.98, k_NTP_ = 50 s^−1^, k_dissociate_ = 1 s^−1^). The rate constants Cumulative function distributions were computed for each transcription start-site distribution.

## Author Contributions

CLT, JOF, and SET cloned, expressed, and purified human core TFIIH. OL completed *in vitro* transcription assays with purified factors. JKR purified human TFIIH and performed kinase assays. JF purified yeast *Sc*TFIIH derivatives and performed *in vitro* transcription assays. SH generated yeast strains for TFIIH core purification. EJT performed ATPase assays, analyzed the data, and did the modeling. EAG, DJT, SET, and SH supervised research and provided funding support. EJT and EAG designed the research and wrote the manuscript with input from all the other authors.

## Acknowledgements

This work was supported by NCI P01 CA092584 (to SET and JOF); F31 CA254478 (to OL); F31 CA250432 (to JKR); NSF MCB-1818147 (to DJT); NIGMS R01GM110387 (to SET); NIGMS R01GM110064 (to DJT); NIGMS 2R01GM053451 (to SH); and NIGMS R01GM120559 (to EAG).

