## Supplemental Figures for "The role of XPB/Ssl2 dsDNA translocation processivity in transcription-start-site scanning"

Affiliations:
[1] Biochemistry and Molecular Biophysics, Washington University School of Medicine, St. Louis, MO 63110, USA
[2] Dept. of Biochemistry, University of Colorado, Boulder, CO 80303, USA.
[3] Division of Basic Sciences, Fred Hutchinson Cancer Research Center, Seattle, WA 98109, USA
[4] Department of Molecular and Cellular Oncology, The University of Texas MD Anderson Cancer Center, Houston, TX, 77030, USA
[5] Molecular Biophysics and Integrated Bioimaging, Lawrence Berkeley National Laboratory, Berkeley, CA, 94720, USA

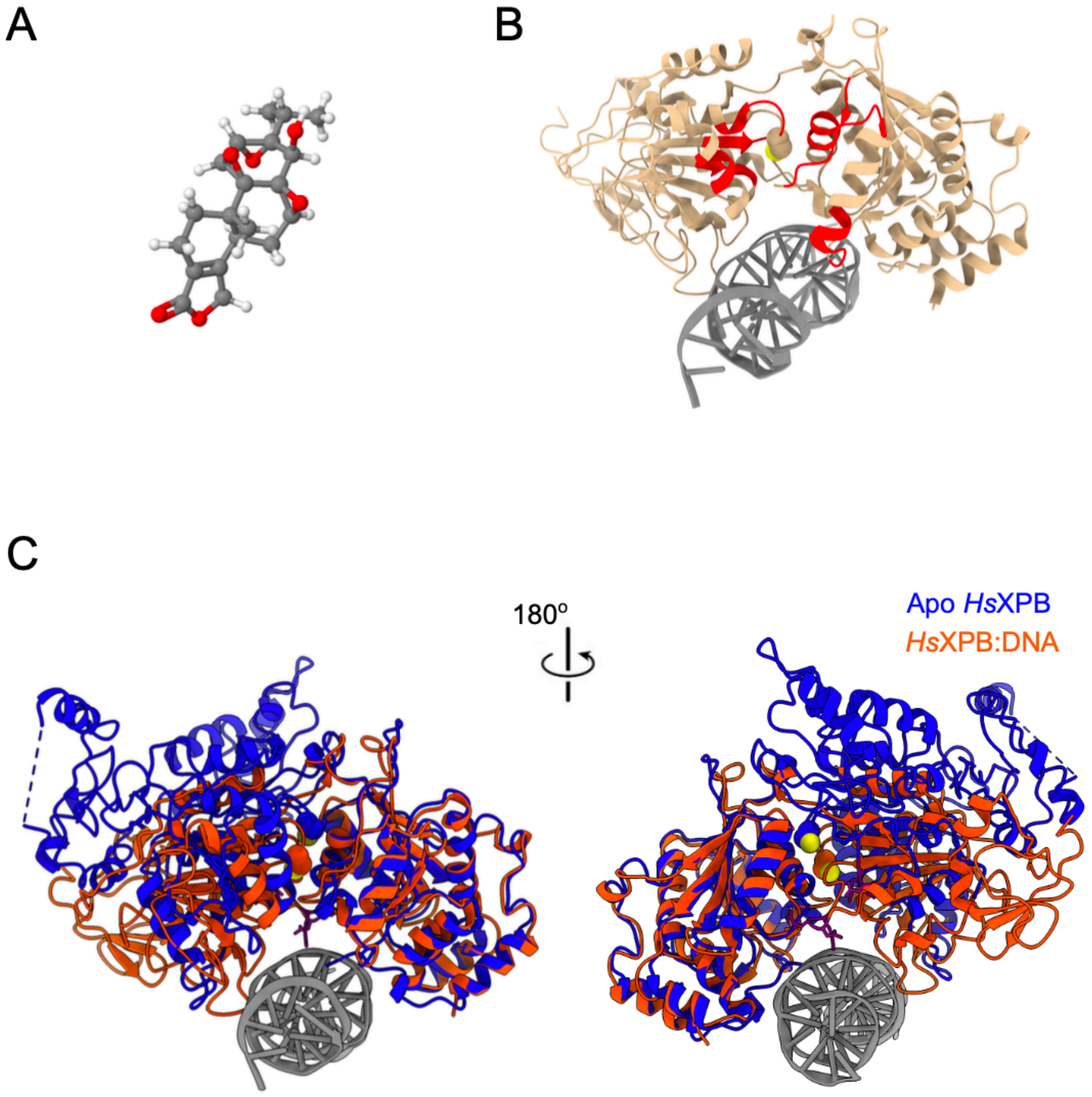


**Figure S1. Site of triptolide modification on XPB potentially disrupts ATP binding site and can be blocked by DNA binding. (A)** Molecular structure of triptolide. **(B)** *Hs*XBP substructure bound to duplex DNA from the *Hs*PIC structure[17] (PDB: 5ivw). Residue C342 of XBP is shown in space fill and is the site of triptolide modification[26]. XBP structure in red are the conserved motifs found in DNA-stimulated ATPases that generally comprise the ATP binding site[28]. **(C)** Alignment of the *Hs*XBP substructure bound to duplex DNA from the *Hs*PIC structure[17] (PDB: 5ivw) with *Hs*XBP substructure from apo, core-*Hs*TFIIH structure[27] (PDB: 6o9m; blue) using Chimera[42]. Residue C342 is shown space filled and becomes less exposed in the DNA bound structure.

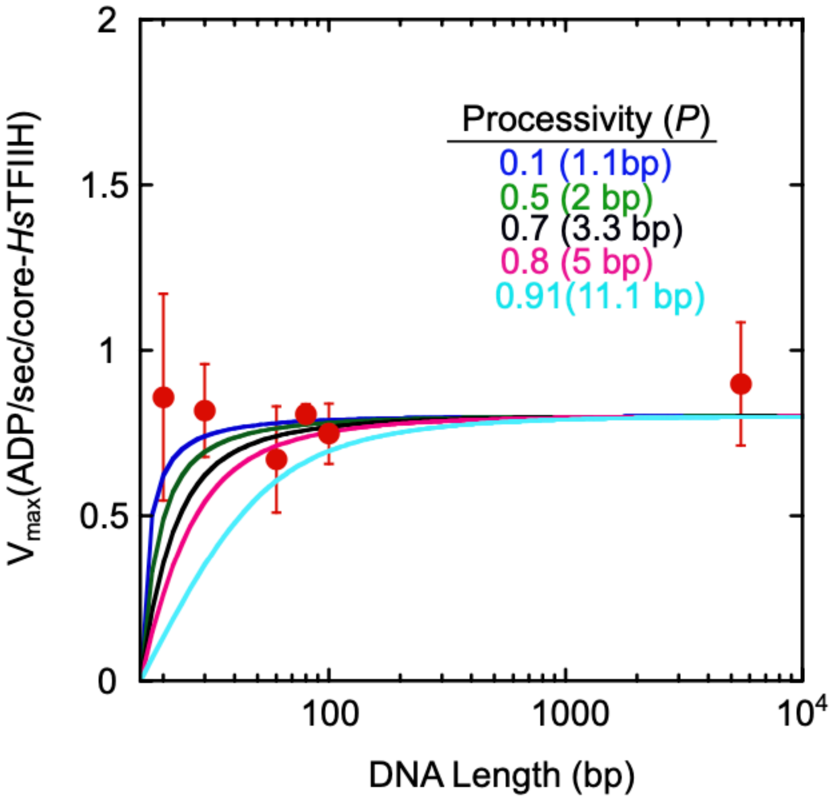


**Figure S2. Estimating an upper limit on *Hs*TFIIH dsDNA translocation processivity.** A series of $V_{max}$ length dependencies calculated from equation (1) using different processivities (P) are plotted along with the observed core-*Hs*TFIIH $V_{max}$ values. The $V_{max}$ length dependencies for P = 0.1 and 0.5 are within the error of all the observed $V_{max}$ values, indicating both are consistent with the data. Thus, an upper limit on the translocation processivity is 0.5 (i.e. ~2 bp translocated on average per binding event).
